# Corticosterone exposure causes long-term changes in DNA methylation, physiology, and breeding decisions in a wild bird

**DOI:** 10.1101/2023.07.07.548150

**Authors:** Conor C. Taff, Sabrina M. McNew, Leonardo Campagna, Maren N. Vitousek

**Affiliations:** CCT, MNV, & LC: Department of Ecology & Evolutionary Biology and Lab of Ornithology, Cornell University; SMM: Department of Ecology & Evolutionary Biology, University of Arizona

**Keywords:** reduced representation bisulfite sequencing, stress, endocrinology, carryover effects, tree swallow

## Abstract

When facing challenges, vertebrates activate an evolutionarily conserved hormonal stress response that can dramatically alter behavior and physiology. Although this response can be costly, conceptual models suggest that it can also recalibrate the stress response system, priming more effective responses to future challenges. Little is known about whether this process occurs in wild animals, particularly in adulthood, and if so, how information about prior experience with stressors is encoded. One potential mechanism is hormonally-mediated changes to DNA methylation. We simulated the spikes in corticosterone that accompany an acute stress response using non-invasive dosing in female tree swallows (*Tachycineta bicolor*) and monitored the phenotypic effects one year later, and DNA methylation both shortly after treatment and a full year later. The year after treatment, experimental females had stronger negative feedback and initiated breeding earlier – traits that are associated with stress resilience and reproductive performance in our population – and higher baseline corticosterone. We also found that natural variation in stress-induced corticosterone predicted patterns of DNA methylation, including methylation of the MC2R gene, which encodes the adrenocorticotropic hormone receptor. Finally, corticosterone treatment causally influenced methylation on short (1-2 weeks) and long (1 year) time scales; however, many of these changes did not have clear links to functional regulation of the stress response. Taken together, our results are consistent with corticosterone-induced priming of future stress resilience, and support DNA methylation as a potential mechanism. Uncovering the mechanisms linking experience with the response to future challenges has implications for understanding the drivers of stress resilience.

**SIGNIFICANCE:** A stress response to an environmental challenge can preserve an individual’s fitness and may even prime them to survive similar challenges in the future. What mechanisms underlie priming is unclear, but epigenetic alterations to stress-related genes are one possibility. We experimentally tested whether increasing corticosterone in free-living swallows had lasting phenotypic or epigenetic effects. A year after treatment, females who received corticosterone had altered stress physiology and bred earlier than control birds, traits that are associated with higher fitness. Treatment also altered DNA methylation and methylation of the MC2R gene was correlated with stress physiology. This study adds to a growing body of literature suggesting that epigenetic changes are key to animals’ response to a changing environment.

## INTRODUCTION

Wild organisms regularly encounter challenging conditions that require rapid behavioral and physiological responses. In vertebrates, the glucocorticoid mediated stress response plays an essential role in allowing animals to successfully avoid or tolerate stressors (1, 2). While an appropriate response is beneficial (2), an inappropriate or prolonged elevation of glucocorticoids can result in a variety of negative consequences for health and fitness (3). Accordingly, the dominant paradigm in behavioral ecology and endocrinology is that the immediate benefits of the stress response are balanced by long-term costs.

But are the long-term effects of a physiological stress response always costly? Some conceptual models propose that activating the stress response system–even in adulthood–primes more effective responses to future challenges (4, 5). These models predict that initiating a response calibrates the stress response system, increasing organismal resilience or robustness. Because physiological priming results from activating the stress response system rather than from learning, it could occur even in the absence of exposure to an identifiable external threat. Physiological studies have provided some evidence that stressor priming can occur, including outside of critical developmental periods (6, 7). However, we know little about the degree to which stressor priming operates in natural populations, affecting later life behavior, physiology, and fitness. Similarly, the mechanism(s) that link activation of the stress response to physiological regulation of subsequent responses are not well understood.

One mechanism that could play a role in the calibration of stress response systems is changes to DNA methylation. Epigenetic modification by DNA methylation can alter phenotypes by making genes or promoters more or less accessible for transcription (8–10). Early life experiences can have profound programming effects on DNA methylation patterns that often persist throughout the individuals’ lifetime (11). For example, classic work in lab rodents demonstrates that early life experiences regulate methylation of the gene producing the glucocorticoid receptor, which results in lifelong changes to glucocorticoid secretion in response to challenges (12, 13). A growing number of studies in wild animals also demonstrate early life programming of DNA methylation patterns in wild populations resulting from dominance hierarchies (14), brood size (15, 16), temperature and weather (17, 18), or landscape features (19).

While they are less well documented, experiences during adulthood can also result in changes to DNA methylation and these adjustments can occur rapidly (17, 20). For example, brief periods of experimental competition and aggression in tree swallows (*Tachycineta bicolor*) resulted in altered DNA methylation of brain regions associated with hormone signaling, suggesting a priming effect in preparation for future aggression (20). Conceptual models of the stress response have long recognized that the sequence, frequency, duration, and intensity of stressors should change the optimal behavioral and physiological response (21, 22). Yet it is often unclear how the experience of challenges during adulthood would be biologically encoded to alter responses to future challenges. Altered DNA methylation is a promising mechanism because i) it can change rapidly even during adulthood, ii) it can persist over moderate to long time scales, iii) it has been shown to change with challenging experiences, and iv) it can directly alter an individual’s phenotype.

We experimentally simulated a series of acute corticosterone spikes using a non-invasive dosing procedure (23) and monitored both long-term phenotypic effects and changes to DNA methylation. In this population we previously found that brief increases in corticosterone have lingering effects on behavior and performance within a breeding season (23, 24) and that genome-wide methylation predicts resilience to experimental challenges (25). Here, we extended those results to ask whether brief increases in corticosterone altered regulation of the stress response and breeding decisions a full year later. We coupled this approach with reduced representation bisulfite sequencing (RRBS) and a newly improved reference genome assembled for this study to examine genome-wide patterns of DNA methylation at high resolution. We used RRBS to first assess covariation between methylation and natural variation in corticosterone regulation during an acute stress response. Next, we compared DNA methylation in corticosterone treated females to controls to determine whether brief increases in corticosterone resulted in altered DNA methylation at either short (1-2 weeks) or long (1 year) timescales.

If activation of the stress response machinery primes future coping ability (4, 5), experimentally treated females should exhibit long-term (across-year) differences in key phenotypic traits. Specifically, we predicted that in the year after treatment females would have a robust stress-induced increase in corticosterone and strong negative feedback, traits that have been previously shown to predict stress resilience in this population (26), and initiate breeding earlier, which is associated with higher reproductive performance in tree swallows (27). Given previous work demonstrating a correlation between coping ability and both genome-wide methylation (25) and natural variation in rapid corticosterone regulation (28), we also predicted that natural variation in corticosterone (baseline, stress-induced increase, and efficacy of negative feedback) would be associated with DNA methylation. However, a correlation here could arise through early life programming, prior activation of the acute corticosterone response, or any conditions that impact the regulation of both methylation and corticosterone (e.g., body condition). In contrast, for the experimental manipulation we predicted that differences in DNA methylation between control and treatment groups would only be present if brief increases in corticosterone have a causal effect on altering methylation patterns. We assessed the time course and persistence of any such changes using comparisons 1-2 weeks after treatments and 1 year after treatments. If methylation changes play a role in altering future corticosterone secretion then we expected to find more differences near genes and promoters associated with endocrine regulation.

## RESULTS

### Corticosterone and breeding timing

Birds that were treated with corticosterone in year one had higher baseline corticosterone in year two (corticosterone treatment *β* = 3.34; 95% confidence interval = 1.43 to 5.24; Figure 2A; Table S2). Prior year treatment was not related to stress-induced corticosterone (Table S2), but females that were previously corticosterone-exposed had lower post-dexamethasone corticosterone, indicating more robust negative feedback (*β* = -4.86; 95% confidence interval = -9.01 to -0.70; Figure 2B; Table S2). Finally, corticosterone-exposed females initiated their nesting attempt earlier in the following year (*β* = -2.34; 95% confidence interval = -4.49 to -0.20; Figure 2C; Table S2).

**Figure 1:**
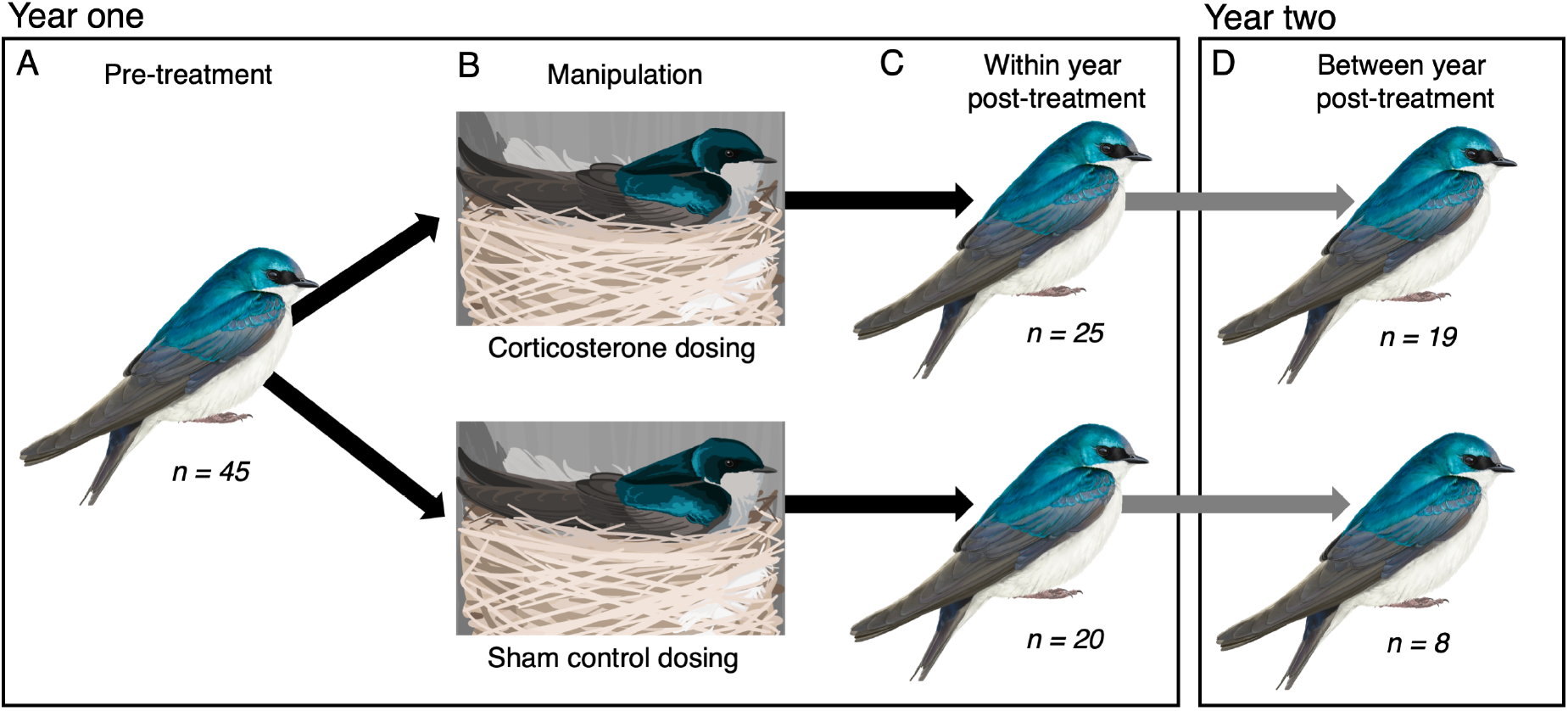
Schematic illustration of the experimental treatment and samples collected for RRBS. Models comparing natural corticosterone to DNA methylation used pre-treatment samples (A). After treatments were applied (B), models testing for within-year effects of treatment used post-treatment samples (C), while controlling for initial methylation (A). Models testing for between-year effects used post-treatment samples (D), while controlling for initial methylation (A). See text for description of birds included in treatment and control groups and additional samples used for analyses not focused on methylation. Note that sample sizes indicate number of samples used in sequencing and analysis, but are not indicative of survival in each treatment.

**Figure 2:**
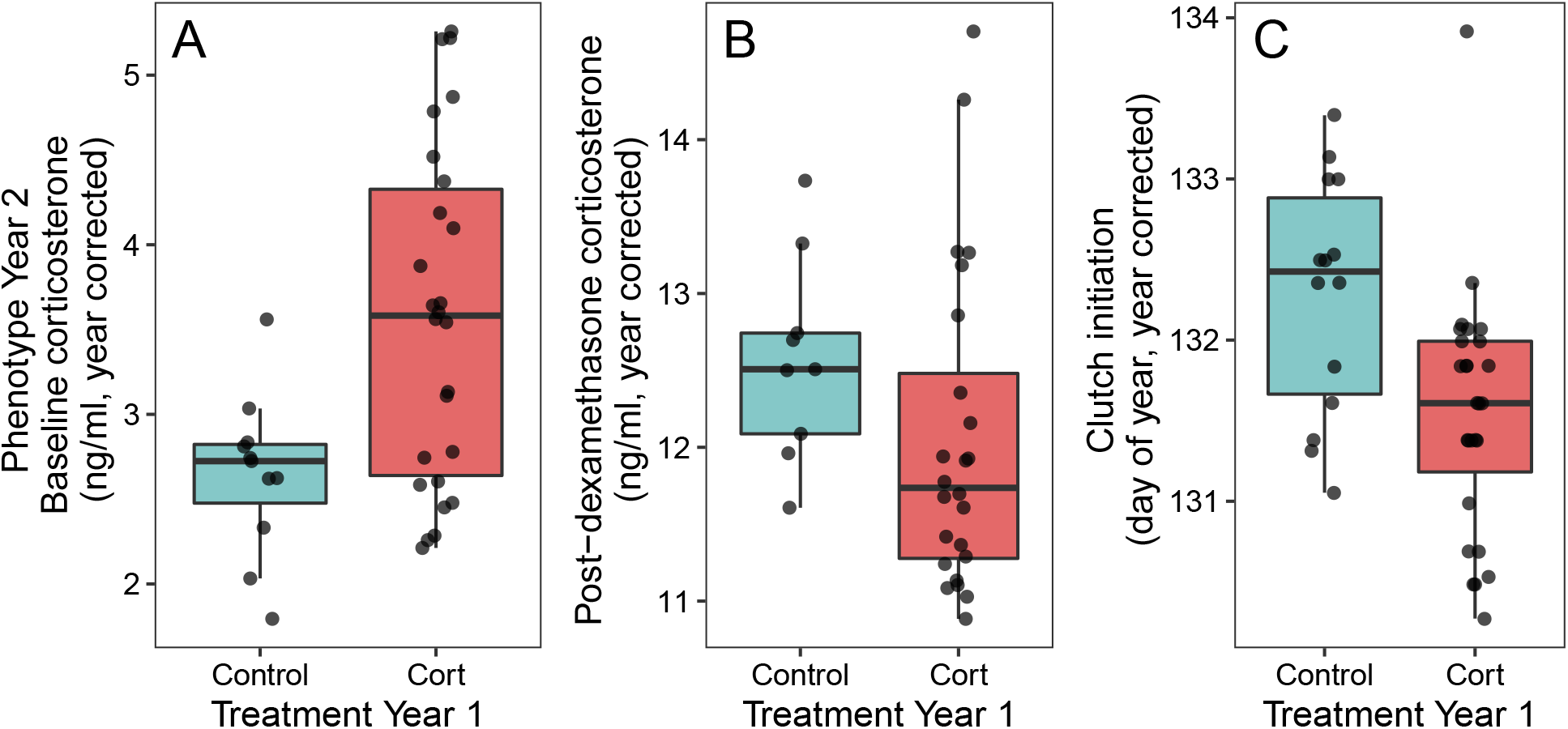
Effect of corticosterone treatment in one year on measures of baseline corticosterone (A), post-dexamethasone corticosterone (B), and clutch initiation date (C) in the following year. Points are partial residuals of raw data collected for average year effects. Boxes and whiskers show the median, interquartile range, and 1.5 times IQR.

### Association between natural or experimental corticosterone and methylation

Using pre-treatment samples, we found that methylation percentage at 116 CpGs out of 78,143 tested was significantly associated with baseline corticosterone after FDR correction (Figure 3A; Table S1). For stress-induced corticosterone, we found a similar association at 356 out of 78,027 CpGs that were tested (Figure 3B; Table S1). For post-dexamethasone injection samples, we found an association between corticosterone and methylation at 735 out of 69,189 CpGs tested (Figure 3C; Table S1).

**Figure 3:**
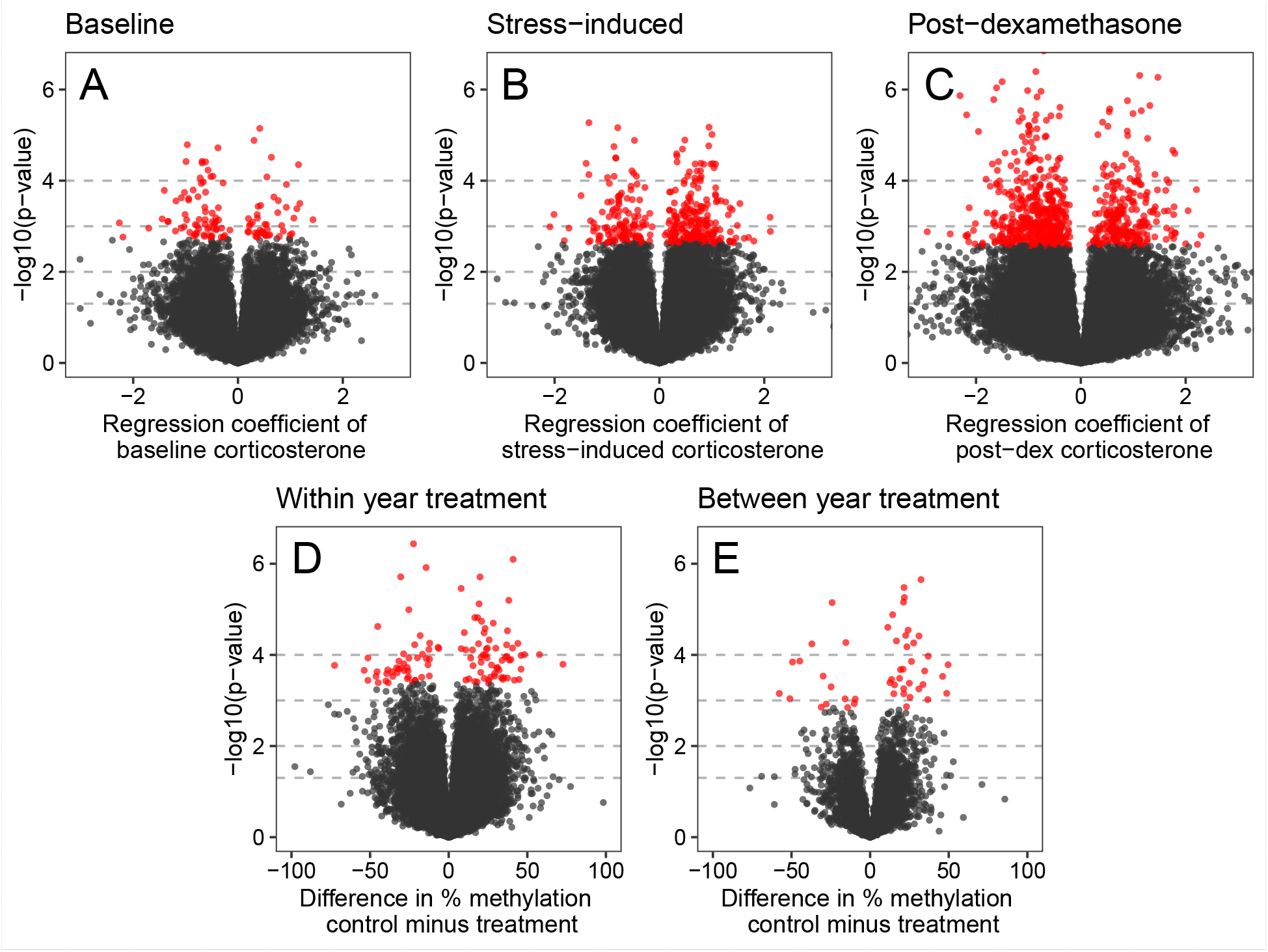
Association between DNA methylation and corticosterone from GLMMs based on observational and experimental study components. Panels A, B, and C show the pre-treatment regression coefficient for baseline corticosterone, stress-induced corticosterone, and post-dexamethasone corticosterone on methylation percentage, respectively. Panels D and E show the difference in methylation for control vs. treatment groups after accounting for pre-treatment methylation percentage for samples 1-2 weeks after treatment (D) and 1 year after treatment (E). In all plots, -log base 10 p-values are shown on the y axis with red points indicating CpGs that were significantly associated with corticosterone after applying the false discovery rate correction. Horizontal dashed lines indicate p-values of 0.05, 0.01, 0.001, and 0.0001 moving from the bottom to top of each plot to aid in interpretation.

In models examining the causal effect of corticosterone treatment, we found that for samples collected within the same breeding season 1-2 weeks after treatment, 111 out of 48,070 CpGs tested showed evidence of differential methylation after FDR correction (Figure 3D; Table S1). We had fewer individuals and fewer CpGs that passed filtering for comparisons one year after treatment, but we found that 49 out of 6,787 CpGs tested were differentially methylated between treatment and control groups after one year (Figure 3E; Table S1). Although we were primarily interested in treatment effects, these models also showed that pre-treatment methylation at a given CpG site generally predicted post-treatment methylation both within a year (Figure S3A) and for samples collected one year later (Figure S3B).

### Association between differentially methylated CpGs and genes

We found that CpGs that were significantly associated with baseline corticosterone, stress-induced corticosterone, and post-dexamethasone corticosterone were located in or near a total of of 32, 176, and 236 identifiable genes, respectively (Table S3). When comparing differentially methylated CpGs after treatment effects, within-year and between-year CpGs were located in or near 52 and 16 genes, respectively (Table S3). A subset of these genes were identified in two or three different comparisons (Figure 4). Because of our filtering process many genes were not tested in each comparison (i.e., the background set of possible genes tested differed for each comparison).

**Figure 4:**
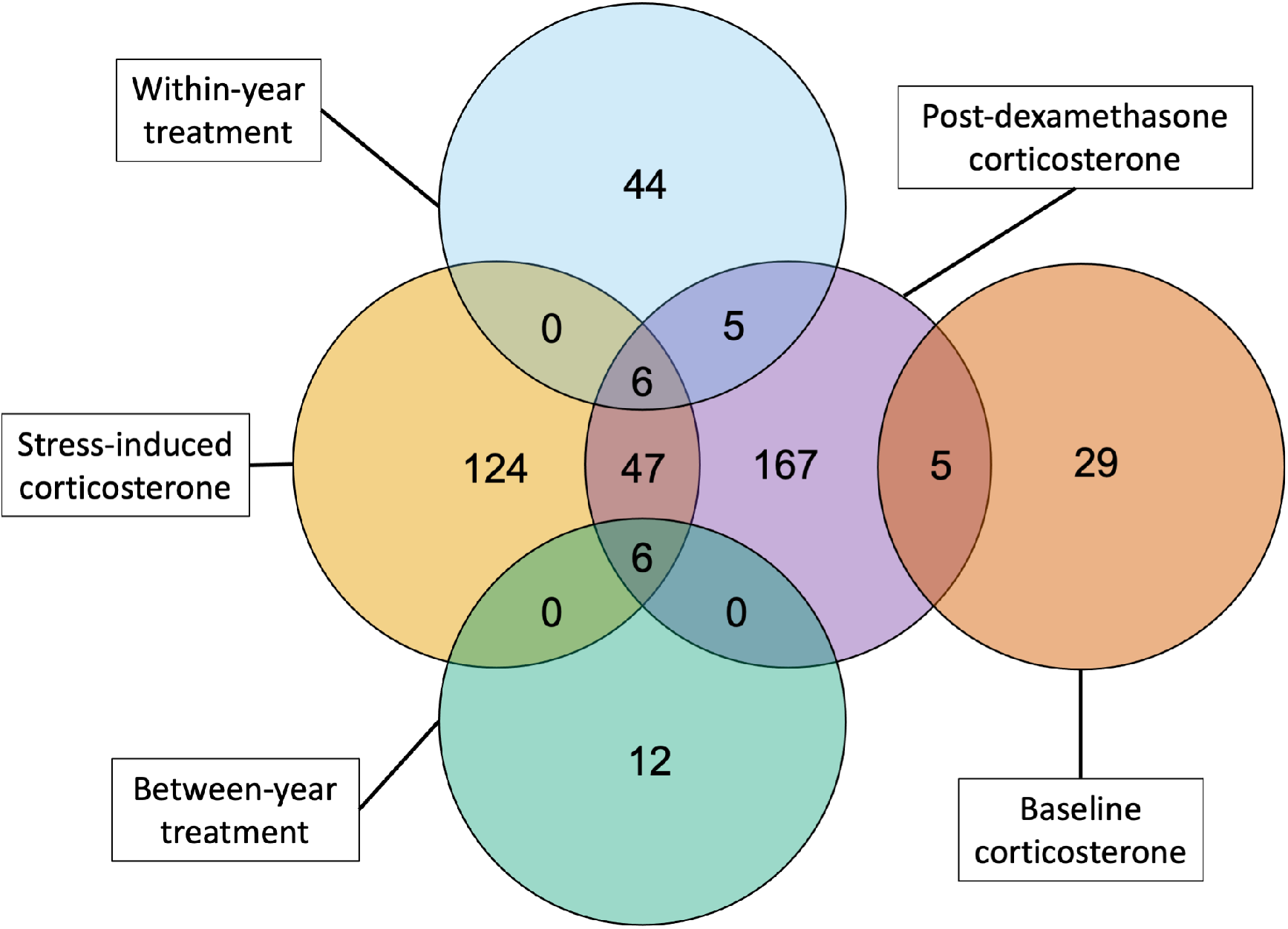
Number of genes near CpGs that were significantly associated with natural variation in corticosterone (baseline, stress-induced, and post-dexamethasone) or with experimental corticosterone elevation (within-year and between-year). No identified genes were shared between the comparisons with circles that do not overlap.

In examining the function of genes identified in this process, only one was obviously directly connected to regulation of the hypothalamic-pituitary-adrenal (HPA) axis. We found that individuals with higher stress-induced corticosterone in the observational dataset had higher methylation at a CpG associated with the MC2R gene, which is responsible for making the ACTH receptor (Figure S4). Several other genes known to be associated with the HPA axis (e.g., CRH, CRHR1, FKBP5) did not have any CpGs near them in the background set, so we could not test for differences associated with these genes.

### GO term analysis

Using the gene lists from Table S3 as input, we identified GO terms that were significantly associated with each comparison. With the false discovery rate set at 0.05, we identified 14 GO terms associated with baseline corticosterone, 22 terms associated with stress-induced corticosterone, 10 terms associated with post-dexamethasone corticosterone, 2 terms for the within-year treatment effect, and 27 terms for the between-year treatment effect (Table S4). None of these lists resulted in any clear clustering of processes using the REVIGO visualization tool and many terms were repetitive and attributable to the same few gene associations.

Baseline corticosterone was associated with photoreceptor activity and response to light, which was primarily driven by opsin and rhodopsin gene associations (OPN1SW, RHO, LWS). Stress-induced corticosterone was associated with a wider range of processes connected to a larger set of genes. These included a variety of cell signaling and receptor pathways, such as the ACTH association described above (MC2R). Post-dexamethasone corticosterone was primarily associated with signaling receptor activity driven by a relatively large number of associated genes (Table S4).

Differentially methylated CpGs for within-year corticosterone treatment were only related to two GO terms associated with structural cell components and attributable to genes of unknown function. Between-year corticosterone treatment was associated with a variety of GO terms having to do primarily with transmembrane receptor signaling, but nearly all of these terms were selected from the same set of gene associations (BMPR1A and B, ACVR1, and TGFBR1).

## DISCUSSION

We found that experimental increases in corticosterone induced long-term phenotypic changes. Females that experienced a few brief spikes in exogenous corticosterone had stronger negative feedback in the HPA axis and bred earlier in the subsequent year; these characteristics are typically associated with high stress resilience and reproductive success in this population. Furthermore, natural variation in corticosterone was correlated with DNA methylation, and experimental treatments altered DNA methylation patterns. Importantly, regulation of DNA methylation in response to corticosterone occurred rapidly in adults (within days) and resulted in detectable changes at least one year after treatment, paralleling the changes in physiological and behavioral phenotypes. Taken together, these results support the idea that the activation of the stress response machinery changes traits associated with stress resilience, and thus may prime future responses to challenges. DNA methylation could act as a key mechanism linking the prior experience of stressors–including during adulthood–to subsequent coping ability. Rapid endocrine flexibility and adaptive calibration of the stress response have emerged as key determinants of resilience to challenges (29–31) and understanding the mechanistic basis of these patterns is an important step in predicting when flexibility is sufficient for coping with changing conditions.

The changes in phenotype that we detected one year after experimentally elevating corticosterone partially matched our predictions if exposure altered phenotype in ways that would increase future stress resilience. We found that, compared to controls, experimental females initiated breeding earlier and had stronger negative feedback in the subsequent year. In tree swallows, clutch initiation date is a strong predictor of both seasonal and lifetime reproductive success and is often considered a proxy for individual quality or condition (27). Similarly, the strength of negative feedback is consistently the best physiological predictor of coping ability and reproductive success both under natural conditions and after imposing experimental challenges (24, 26). However, contrary to our prediction, treatment had no effect on stress-induced corticosterone the following year. We also found that corticosterone dosed females had higher baseline corticosterone and no difference in stress-induced corticosterone one year after treatment. Baseline corticosterone does not predict stress resilience in this population (26). However, because baseline corticosterone often increases in preparation for periods of high energetic demands, including the demands of reproduction (the cort-adaptation hypothesis, 32, 33, 34), these results might reflect an increased allocation to breeding in subsequent years in corticosterone treated females. For example, female European starlings (*Sturnus vulgaris*) manipulated to increase parental investment increased their baseline corticosterone during incubation (35). Similarly, tree swallows that increase baseline corticosterone more over the reproductive period provision offspring at higher rates (32). Thus, our results might represent a combination of long-term priming effects coupled with the immediate energetic demands of breeding earlier.

Our study also adds to the growing recognition of bidirectional links between coping ability and DNA methylation. While this relationship has been demonstrated in laboratory-based model systems (12, 13) the potential for environmental stressors to trigger methylation, and affect subsequent coping ability, has only recently been explored in wild animals. Early results in wild animals suggest patterns similar to those seen in laboratory rodents. For example, early life maternal care and social connections in spotted hyenas (*Crocuta crocuta*) predict DNA methylation and glucocorticoid regulation as an adult (14, 36). Similar effects can play out in adulthood; for example, in savannah baboons (*Papio cynocephalus*) high social status as an adult is associated with more rapid changes to DNA methylation (epigenetic aging) as a consequence of the social stress that accompanies high status (37). Our results are consistent with the results derived from lab rodents, wild mammals, and a growing number of studies in wild birds (18, 38), suggesting that flexible adjustment of methylation may be a general mechanism by which prior experiences of stressors are encoded in order to modulate future responses to challenges.

While there has been a rapid increase in studies of methylation in wild birds in recent years (18, 19, 39), relatively few studies have sampled the same adults multiple times. Our study design allowed us to assess the stability of genome-wide DNA methylation within individuals. We found that many CpGs that we interrogated had large between individual differences in methylation and that those differences were typically stable even in samples collected one year apart. Compared to these individual differences, flexible changes in methylation were relatively smaller and detectable at fewer CpGs. The stable individual differences that we detected might represent the consequences of early life conditions (14–16). For example, early life climate conditions are related to lifelong methylation of the glucocorticoid receptor gene in superb starlings (*Lamprotornis superbus*, 18). We could not assess the possibility of a similar pattern in our study because we did not have any information on early life conditions for our birds. Regardless of the source of these initial differences, our results clearly demonstrate that detecting subtle adjustments of methylation in adulthood to any treatment of interest will often require accounting for pre-treatment methylation.

The functional consequences of most of the specific methylation changes that we detected are somewhat unclear. We did find that stress-induced corticosterone was correlated with methylation of a CpG associated with the MC2R gene, which encodes the ACTH receptor. Individuals with a more robust stress response had higher methylation at this CpG. Higher methylation is expected to be associated with lower gene expression (8, 10), suggesting that individuals with a more robust corticosterone response might have lower ACTH receptor expression. It isn’t clear why lower ACTH receptor expression would result in higher stress-induced corticosterone, but because regulation of the HPA axis can occur at multiple levels with bi-directional feedback, the result may instead reflect a downregulation of ACTH receptor expression in response to robust activation of other components of the HPA axis. Pairing RRBS with gene expression measurements and comparisons in different tissues would be helpful to understand these patterns (e.g., 40).

In contrast to stress-induced corticosterone, none of the genes or GO terms associated with natural variation in baseline corticosterone, post-dexamethasone corticosterone, or with treatments had clear connections to HPA axis regulation. We previously found that non-specific, genome-wide methylation predicts stress resilience to experimental challenges in this population (25). Thus, the differences that we detected might reflect large-scale regulation of methylation rather than targeted regulation of sites with specific functional consequences. Alternatively, some of the changes that we detected might have functional effects on stress response calibration that are not obvious from the known effects of those genes. In support of this idea, we did find some overlap between methylation in the genes associated with natural variation in corticosterone and with the consequences of our experimental manipulation of corticosterone. In particular, post-dexamethasone corticosterone, which is a strong predictor of stress resilience in tree swallows (24, 26), had the most extensive correlations between methylation and identified genes and some of these genes were shared with the other corticosterone measures and with treatment effects.

Another potential reason for our failure to find clear links between changes in DNA methylation and genes associated with the stress response may result from limitations of our approach. An advantage of RRBS is that it does not rely on pre-selecting candidate genes, but a disadvantage is that not all relevant genes are necessarily tested. After filtering our data, many of the genes with known roles in the HPA axis were not included in comparisons or had coverage at only a few CpG sites. Thus we did not directly test for methylation differences for many key genes. It is possible that deeper sequencing of our libraries would have improved our ability to detect functional differences. Moreover, although we used the most complete reference genome available for tree swallows, many CpGs mapped to predicted genes whos function is unknown. Continued improvement of assembly and annotation for reference genomes of non-model organisms is important for understanding the functional importance of epigenetic changes. Studying DNA methylation in non-model systems is a rapidly developing field and many recent papers outline the pros and cons of various approaches (9, 41, 42). One particularly promising approach that may strike a balance between a focus on candidate genes and the ability to detect genome-wide associations is to combine RRBS with probes that enrich sequences at a potentially large number of target genes (target-enriched enzymatic methyl sequencing, 43).

Regardless of the functional consequences of the changes we detected, we found that brief increases in corticosterone have effects on subsequent corticosterone regulation, breeding decisions, and methylation a full year after dosing ended. At least some of the phenotypic changes we detected support a hormone-mediated priming effect in which activation of the stress response machinery improves the capacity to cope with future challenges, increasing stress resilience. The fact that these changes in phenotype are coupled with changes in methylation patterns implicates the regulation of DNA methylation as a potential mechanism of flexibly adjusting the stress response system based on prior experiences. Understanding the mechanisms that integrate experience with future stress responsiveness has important consequences for predicting how and when individuals can cope with repeated exposure to challenges. Conceptual models of the stress response suggest that while repeated challenges can sometimes impose long-term costs (21, 44), activation of the stress response at other times may prime more effective responses to future challenges (4, 45). Studying the mechanisms by which stress exposure is encoded biologically will help to differentiate these possibilities and shed light on when and how individuals succeed or fail through flexible regulation of the physiological response to challenges.

## MATERIALS AND METHODS

We studied tree swallows breeding at field sites in and around Ithaca, New York, U.S.A. from April to July 2014 to 2017. This population of tree swallows has been continuously studied since 1986 and we followed well-established monitoring protocols (for details see 27). Adult females were captured on day 6 to 7 after the beginning of incubation and again on day 3 to 7 after eggs had hatched. In the year after treatment, any returning females were captured on day 6 to 7 of incubation. At each capture we collected blood samples (< 70*µ*l each) to measure baseline (< 3 minutes) and stress-induced (30 minutes) corticosterone (23). Immediately after the second blood sample was taken, females were injected with 4.5 *µ*l/g of dexamethasone in the pectoralis muscle, which stimulates strong negative feedback (Mylan 4mg ml_-1_ dexamethasone sodium phosphate; previously validated in 26). A final blood sample was collected 30 minutes after injection to measure the efficacy of negative feedback. We also collected a set of standardized morphological measurements and monitored reproductive success (23). All birds received a unique USGS aluminum band and passive integrated transponder (PIT) tag if they were not previously banded.

Between the first and second capture in year one, females were randomly assigned to either a control or experimental treatment group (experiment schematic and sample sizes at each stage are shown in Figure 1). In the experimental group, we simulated a brief spike in corticosterone once per day on five consecutive days between the two captures. To accomplish this, we applied two 60 *µ*l doses of corticosterone dissolved in DMSO gel one hour apart to a fake egg anchored in the nest cup at a randomly chosen time during the day when females were absent from the nest. Upon returning, females incubated the clutch and absorbed the corticosterone across the skin on their brood patch. For the purposes of this study, we considered females as part of the corticosterone treatment group if they received any of the three dose concentrations described in Vitousek et al. 2018 (high = 4 mg ml ^-1^ corticosterone once plus sham once per day; low = 2 mg ml ^-1^ once plus sham once per day; long = 2 mg ml ^-1^ twice per day).

We previously validated that this dosing method results in a brief (< 180 minutes) increase in corticosterone within the range of natural acute corticosterone responses (23). Control nests received either no manipulation or a sham control in which they were dosed as described above but with DMSO gel only with no corticosterone added. We previously found no difference in physiology, behavior, reproductive success, or survival between control and sham control birds receiving this treatment (23, 24) and we combined both control groups in the analyses described here.

For methylation analyses, we focused on the set of females that were manipulated in 2015 and–if they returned–recaptured in 2016. In analyses focused on between year effects of treatments on later corticosterone regulation and breeding decisions, we also included a smaller number of females observed from 2014 to 2015 and from 2016 to 2017. These additional samples included slight variants on corticosterone dosing that we considered part of the corticosterone treatment group (six doses or three doses of 4 mg ml^-1^ corticosterone once per day during incubation, 24). In the year after exposure, we only considered potential carryover effects of treatment on corticosterone regulation at the first capture and on the timing of clutch initiation because some females were subsequently entered into unrelated experiments that could have influenced later season measures.

### Tree swallow reference genome assembly

For this study, we improved upon a previously published reference genome sequenced from a female belonging to this study population (25) by first extracting high molecular weight DNA from this same individual. We performed a phenol-chloroform extraction followed by an ethanol precipitation and finally a bead cleanup. The Duke Center for Genomic and Computational Biology core facility used the DNA to produce a large insert library (15 to 20 kb), which was subsequently sequenced on 3 cells of a Pacific Biosciences RSII instrument. This produced a total of 9.6 Gbp of data with an average read length of 12,053 bp and an N50 subread length of 15,643 bp. We used bamtools version 2.5.1 (46) to merge the reads from the difference cells and retain only those that were longer than 4,500 bp (47.6% of the original raw reads). We improved our first assembly with the PBJelly2 module of PBSuite version 15.8.24 (47), which uses long reads to fill or reduce gaps. This pipeline produced an assembly which was moderately improved from the previous version (25). The total length of the assembly was 1.22 Gb (previously 1.14 Gb) and was contained in 49,278 scaffolds (previously 92,148), with an N50 of 82.9 kb (originally 34 kb) and 1.9% Ns (vs. 5.8%). We assessed the completeness of our assembly by running BUSCO version 5.2.2 (48), using the passeriformes dataset of 10,844 conserved genes. We found 80.5% of these genes in a single and complete copy, 3.9% were fragmented, 1.8% were duplicated, and 13.8% were missing. Finally, we annotated the genome following the pipeline described in Taff et al. (25). The assembly generated for this project is deposited on GenBank (BioProject ID PRJNA553513).

### Sample processing

Blood samples collected in the field were immediately stored on ice in a cooler and processed in the lab within 3 hours of capture. Red blood cells were separated from plasma by centrifugation and added to 1 mL of ice cold cryopreservation buffer (90% newborn calf serum, 10% DMSO, 49). Samples were then frozen at a constant cooling rate with isopropyl alcohol and stored at -80*^◦^* C until further processing. Cryopreserved blood samples were thawed and DNA was extracted using the DNeasy Blood & Tissue spin column extraction kits according to the manufacturer’s protocol (Qiagen Sciences Incorporated).

### Corticosterone and breeding timing data analysis

We used general linear models to compare corticosterone and the timing of breeding between control and experimental females one year after corticosterone manipulations. We fit four models in total with either the date of clutch initiation or circulating corticosterone levels (baseline, stress-induced, or post-dexamethasone injection) as the response variable. Predictors included treatment and year as a categorical fixed effect. The model for stress-induced corticosterone also included baseline corticosterone as a predictor and the model for post-dexamethasone corticosterone also included stress-induced corticosterone as a predictor.

### Reduced representation bisulfite sequencing

We prepared our samples for reduced representation bisulfite sequencing (RRBS) using the Diagenode Premium RRBS Kit and closely following the manufacturer’s protocol (50). Briefly, samples were diluted to 3.85 ng/*µ*l and 26 *µ*l of diluted sample was used for library preparation. The process included enzymatic digestion with Mspl and size selection to increase coverage of CpG-rich regions, such as CpG islands and enhancers. Individual samples received one of twenty-four unique barcodes and were pooled in randomized groups of 8 before bisulfite conversion. We also included a methylated and unmethylated spike in control with each sample to confirm the efficiency of bisulfite conversion.

From the available samples, we selected 120 samples to process from 61 unique birds (three samples per bird n = 14, two samples n = 31, one sample n = 16). Prior to RRBS processing, these 120 samples were randomly sorted to account for any batch effects. Libraries were prepared with the Diagenode kit in two batches (one set of 24 and one of 96). Prepared libraries were checked for the expected size distribution by digital PCR prior to sequencing. Sequencing was performed at the Cornell BRC using NextSeq 1x75 with 20% PhiX and 85% of the normal cluster density. In total, we ran our samples on five sequencing lanes with 24 samples per lane.

Raw sequence data were first processed with Trim Galore! using the default RRBS settings. Visual inspection of FastQC files confirmed high quality reads for all samples. Next, we used Bismark to align each sequence to the prepared genome and extract the methylation status for each CpG, CpH, or CHH site (51). As expected, global methylation at CpH and CHH sites was extremely low (1.0% and 0.6%, respectively, Figure S1) and we only considered methylation at CpG sites in our subsequent analyses. We also used Bismark to determine the methylation conversion efficiency for each sample based on methylated and unmethylated spike in controls and following the instructions in the Diagenode RRBS kit (50, 51).

### General methylation patterns

Our process resulted in 9.8 *±* 4.3 million (SD) total reads per sample (Figure S1). Across all samples, we were able to align 51.1% of the total reads produced, which is comparable to several recent studies in wild birds (39, 52). Spiked controls in each sample indicated that our bisulfite conversion worked efficiently and within the recommended kit parameters (conversion of methylated control sites = 1.9% *±* 1.4; conversion of unmethylated control sites = 99.5% *±* 0.6).

Among 45 pre-treatment samples, we had sufficient coverage to estimate methylation at 148,167 CpGs. In total, the average percentage methylation across all sites was 35.5% *±* 34.0 with a wide distribution (Figure S2A). After assigning CpGs hierarchically to promoter (within 2kb upstream of a TSS) > exon > intron, we found that 12.1% of sites were in promoters, 7.9% in exons, 11.8% in introns, and 68.1% in intergenic regions. At the level of genomic features, promoters had the lowest methylation (median = 5.3%, mean *±* SEM = 20.5% *±* 0.5), introns had intermediate methylation (median = 43.5%, mean *±* SEM = 41.0% *±* 0.5), and exons had the highest methylation (median = 54.3%, mean *±* SEM = 46.7% *±* 0.7). However, each of these features had a wide distribution of methylation percentages across different genes (Figure S2B).

### RRBS Data Analysis

Output data from the sequence processing described above was analyzed in R version 4.1.1 (53). We processed the aligned sequence data with MethylKit (54). Using MethylKit, we extracted the number of total aligned reads and number of methylated or unmethylated reads for each CpG site.

For analyses of corticosterone and treatment associations, we filtered these CpGs to include only those that met the following criteria. First, we required a minimum coverage of 10 reads per sample to retain data for that sample at a given CpG. We further filtered the dataset to remove any CpGs that were mostly invariant (i.e., more than half of samples had methylation percentage of 0 or 100%) as well as CpGs that had extremely low variation (SD less than 5% across all samples, 55, 56). For models comparing treatment effects, we required that females have data at a given CpG from both pre- and post-treatment sampling points to be included. For basic descriptions of methylation patterns, we used all CpGs that had 10 reads or more in the pre-treatment samples.

The built-in differential methylation techniques in MethylKit are designed for two group comparisons with limited flexibility in modeling options. Because we had repeated measures before and after treatments for both groups, we could not specify the necessary models within MethylKit itself. Therefore, we exported and combined the filtered CpG records for all groups so that we could fit generalized linear mixed models (GLMMs) for each CpG site (as in 38) using the glmer function in R package lme4 (57). We fit a separate set of models for natural corticosterone variation (baseline, stress-induced, or post-dexamethasone), within-year treatments, and between-year treatments. Each of these datasets were constructed separately since they included different subsets of both individual birds and of CpGs that met the criteria described above.

For natural variation in corticosterone, we included only the pre-treatment samples. Using these samples, we fit a GLMM for each CpG with the number of methylated and unmethylated reads as the binomial response variable. We fit this set of models separately with baseline, stress-induced, or post-dexamethasone corticosterone as the single continuous predictor variable. The models included a random effect for female identity to account for repeated sequencing of the same CpG sites within each female. We excluded the results for any models that failed to converge because we could not reliably estimate effects in those cases.

For within-year and between-year comparisons after treatments, we fit a single GLMM for each included CpG with the number of methylated and unmethylated reads as the binomial response variable. Predictors included pre-treatment methylation percentage at the CpG being modeled, a fixed effect of treatment (control vs. corticosterone), and a random effect for female identity. In each model, significance of the comparison between control and corticosterone treated birds was assessed using the emmeans package in R (58). We also evaluated the stability of methylation within individuals in these models by summarizing the regression coefficient of pre-treatment methylation on post-treatment methylation.

We accounted for multiple comparisons in each of these analyses by adjusting all p-values using the q-value approach implemented by the qvalue package in R with the false discovery rate set at 0.05 (59). We only report and interpret estimates with q-values < 0.05.

### Annotation of differentially methylated CpGs

After identifying CpGs that were significantly associated with either natural corticosterone or experimental treatment with corticosterone, we identified genes associated with each CpG. We used the bedtoolsr package to select genes that had a significant CpG either within the gene body or within 2 kb upstream of the transcription start site (60). We generated separate lists of genes associated with CpGs for baseline corticosterone, stress-induced corticosterone, post-dexamethasone corticosterone, within-year treatment effects, and between-year treatment effects. For each of these comparisons we also generated a complete list of genes associated with all of the CpGs that passed the filtering criteria described above to be used as a null background list (see below).

Starting with the list of genes associated with each comparison set, we used the DAVID functional annotation tool (61) to test whether our genes were enriched in any molecular functions or biological processes in the Gene Ontology knowledgebase (62, 63). For each comparison we used the custom background list generated above. This background list is important for interpretation because we were only able to test CpGs near a subset of genes in each comparison (number of genes included in testing for baseline corticosterone = 4,143; stress-induced corticosterone = 4,146; post-dexamethasone corticosterone = 3,863; within-year treatment = 2,913; between-year treatment = 452).

Using DAVID we identified a set of GO terms associated with biological processes or molecular functions that were over represented in the list of significant CpGs compared to the background list for that comparison (63). We filtered this list to include only GO terms with p-values < 0.05 after applying a false discovery rate correction. We initially visualized the GO terms for each comparison using REVIGO (64); however, our study identified a relatively small number of GO terms and no clearly identifiable clusters of terms were identified in REVIGO. Therefore, we report the complete list of genes and GO terms associated with CpGs in each comparison.

### Data and code availability

The complete set of bioinformatic processing scripts, R code, and data associated with each sample is available and permanently archived on Zenodo (https://doi.org/10.5281/zenodo.8125153). Raw sequence data from RRBS is available on GenBank (BioProject ID PRJNA953597).

## ACKNOWLEDGMENTS

We thank the members of the tree swallow research team for assistance with fieldwork including, Joseph Byington, Joe Colcombe, Collin Dickerson, Garret Levesque, Jacob Kolenda, Lyra Liu, Sophie Nicolich-Henkin, Teresa Pegan, Avram Pinals, Alyssa Rodriguez, Vanesa Rodriguez-Arcila, Tom Ryan, David Scheck, Lauren Smith, Jocelyn Stedman, and Cedric Zimmer. Bronwyn Butcher and the Lovette Lab provided input on lab methods and members of the Vitousek Lab provided feedback on earlier versions of the manuscript. Charlotte Holden produced the tree swallow illustrations.

## AUTHOR CONTRIBUTIONS

CCT and MNV conducted the field based data collection. CCT and MNV conceived the study. CCT and SMM conducted the lab work for RRBS. CCT and LC conducted the lab work for creating the reference genome and LC carried out the bioinformatics for genome assembly and annotation. CCT analyzed and visualized the data with assistance from SMM and LC. CCT drafted the paper with input from all authors.

## ETHICAL NOTE

All methods were approved by Cornell IACUC and sampling was conducted with appropriate state and federal permits.

## SUPPLEMENTARY FIGURES AND TABLES

**Figure S1:**
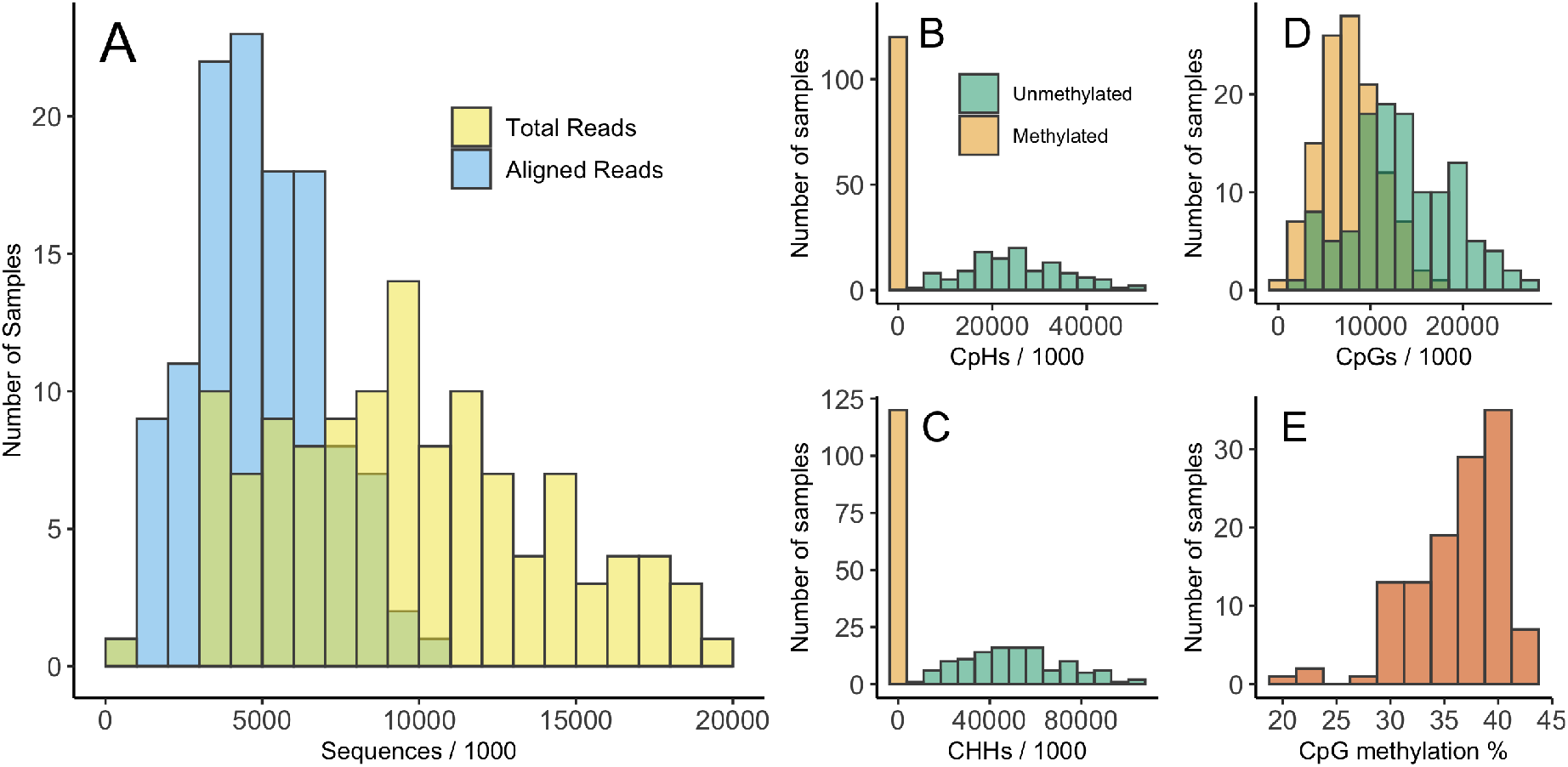
Summary of sequencing and methylation call results from raw sequence data. Panel A shows the distribution of the total number of sequences for each sample and number of sequences that aligned to the tree swallow genome. Panel B shows the number of CpH sites that were methylated or unmethylated for each sample. Panel C shows the number of CHH sites that were methylated or unmethylated for each sample. Panel D shows the number of CpG sites that were methylated or unmethylated for each sample. Panel E shows the percentage of total CpG reads that were methylated by sample. Note that these histograms are based on raw sequencing results that do not account for differential coverage between samples or locations in the genome and are included for illustration only.

**Figure S2:**
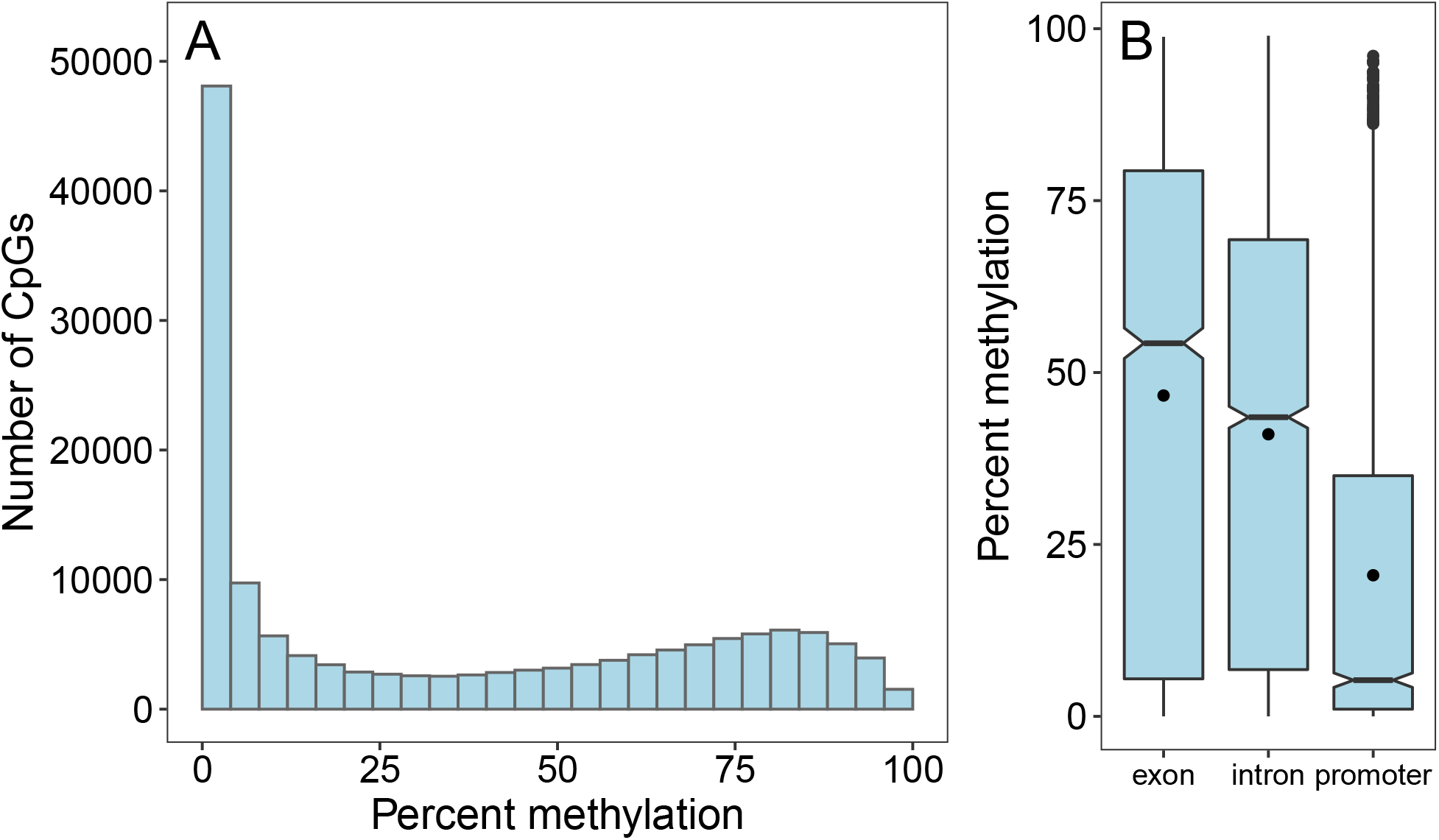
Distribution of methylation percentage across all CpGs and for different genomic features. Panel A shows entire distribution of methylation percentage for all 148,167 CpGs from pre-treatment samples before any filtering. Panel B shows the methylation percentage for exons, introns, and promoters that had CpGs identified within them. Horizontal lines, boxes, and whiskers show the median, interquartile range, and 1.5 times IQR respectively. The black circle within each box is the mean for that feature.

**Figure S3:**
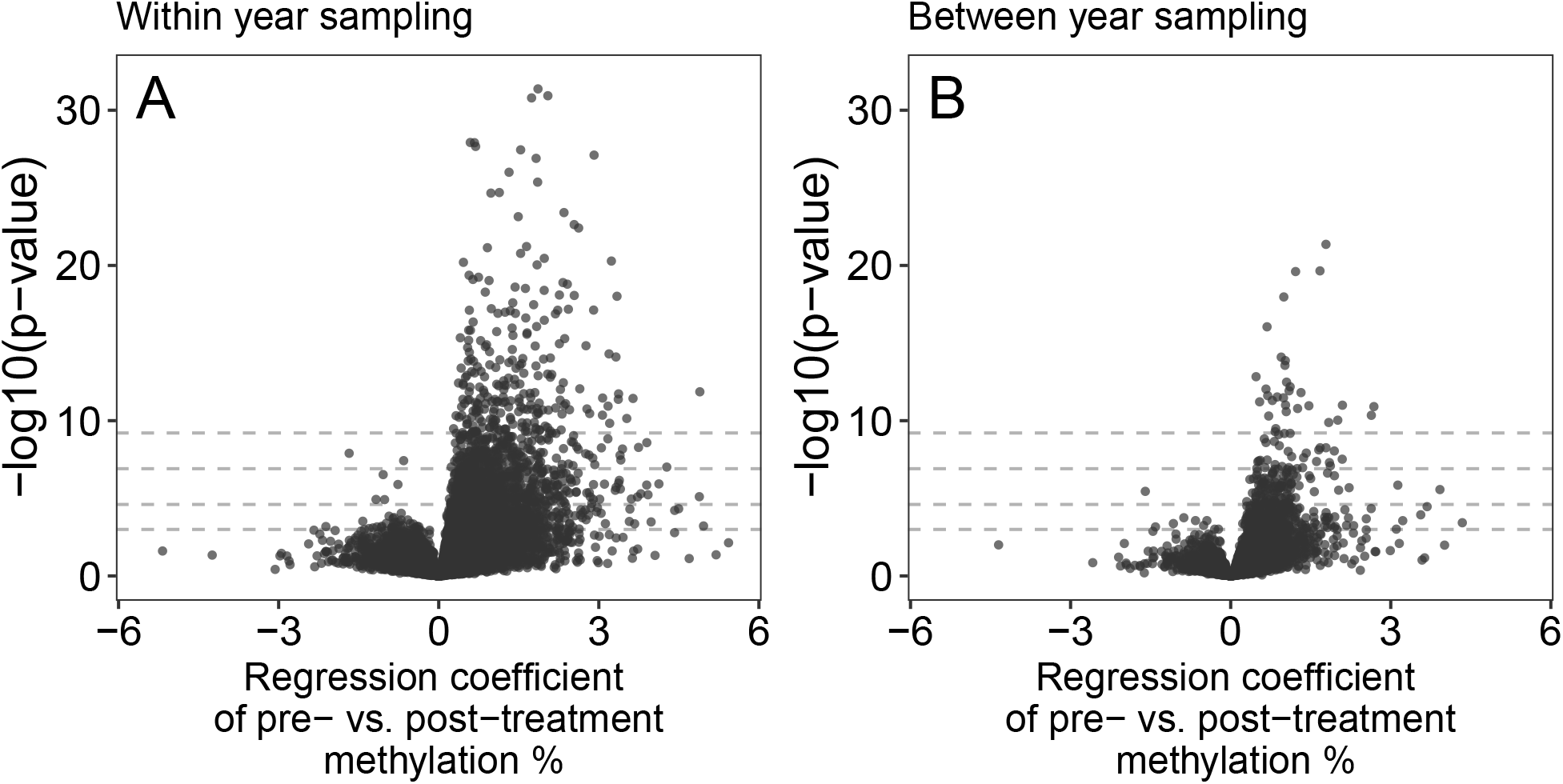
Association between pre-treatment methylation percentage and post-treatment methylation at each CpG for samples collected within a breeding season (panel A) and for samples collected one year apart (panel B). To help with interpretation, horizontal dashed lines indicate p-values of 0.05, 0.01, 0.001, and 0.0001 moving from the bottom to top of the plots.

**Figure S4:**
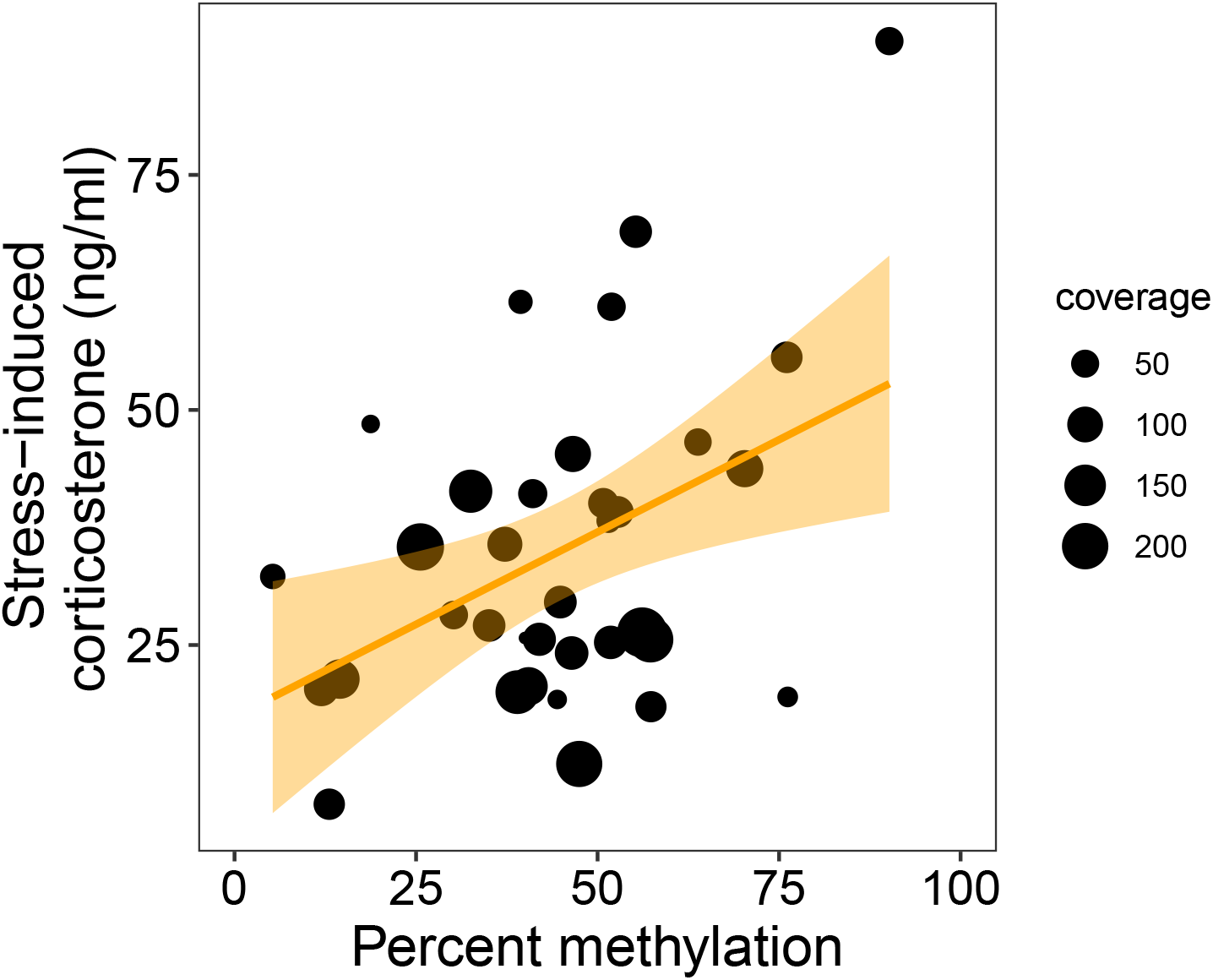
Percent methylation in relation to stress-induced corticosterone at the CpG near the MC2R gene that was significantly associated with corticosterone. Size of circles indicates sequence coverage for each sample. The trendline and confidence interval are shown for illustration but significance was assessed using the binomial GLMM described in the text.

**Table S1:**
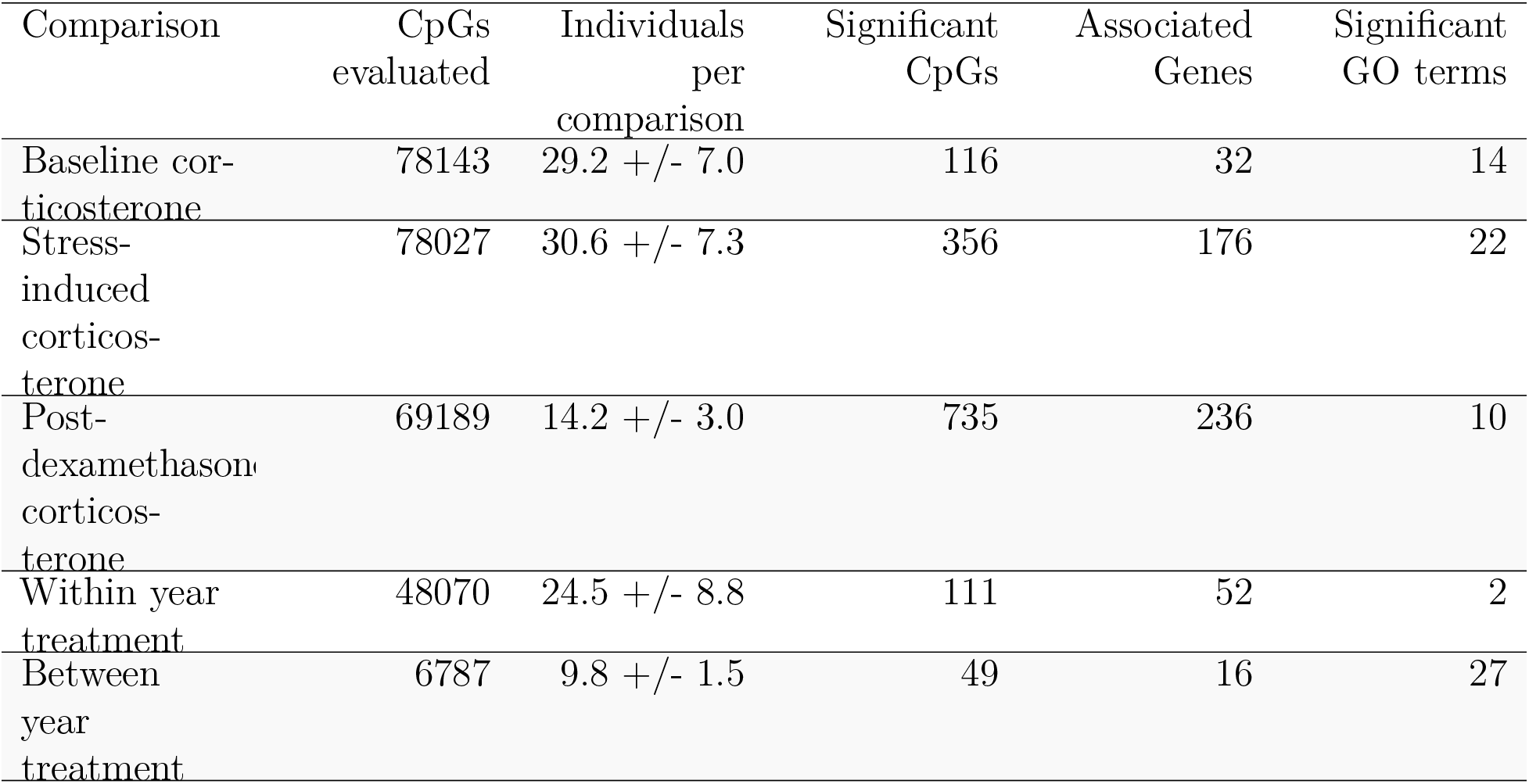
Summary of GLMMs for each comparison with the number of CpGs significantly correlated with corticosterone or differentially methylated between treatment groups. One model was fit for each CpG site; see text for description of models.

**Table S2:**
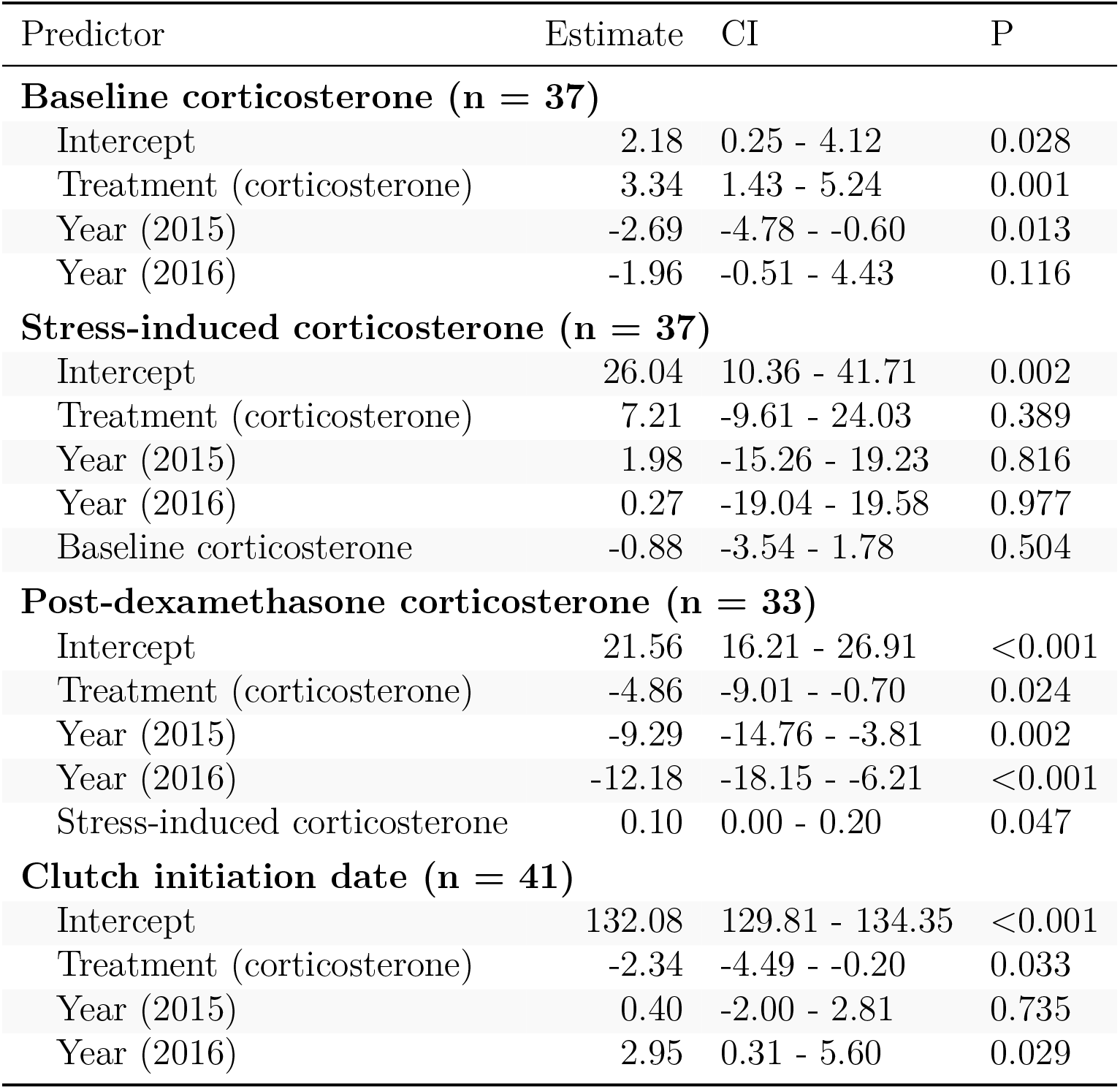
Results of GLMs for corticosterone and clutch initiation date in the year after experimental treatments were applied.

**Table S3:**
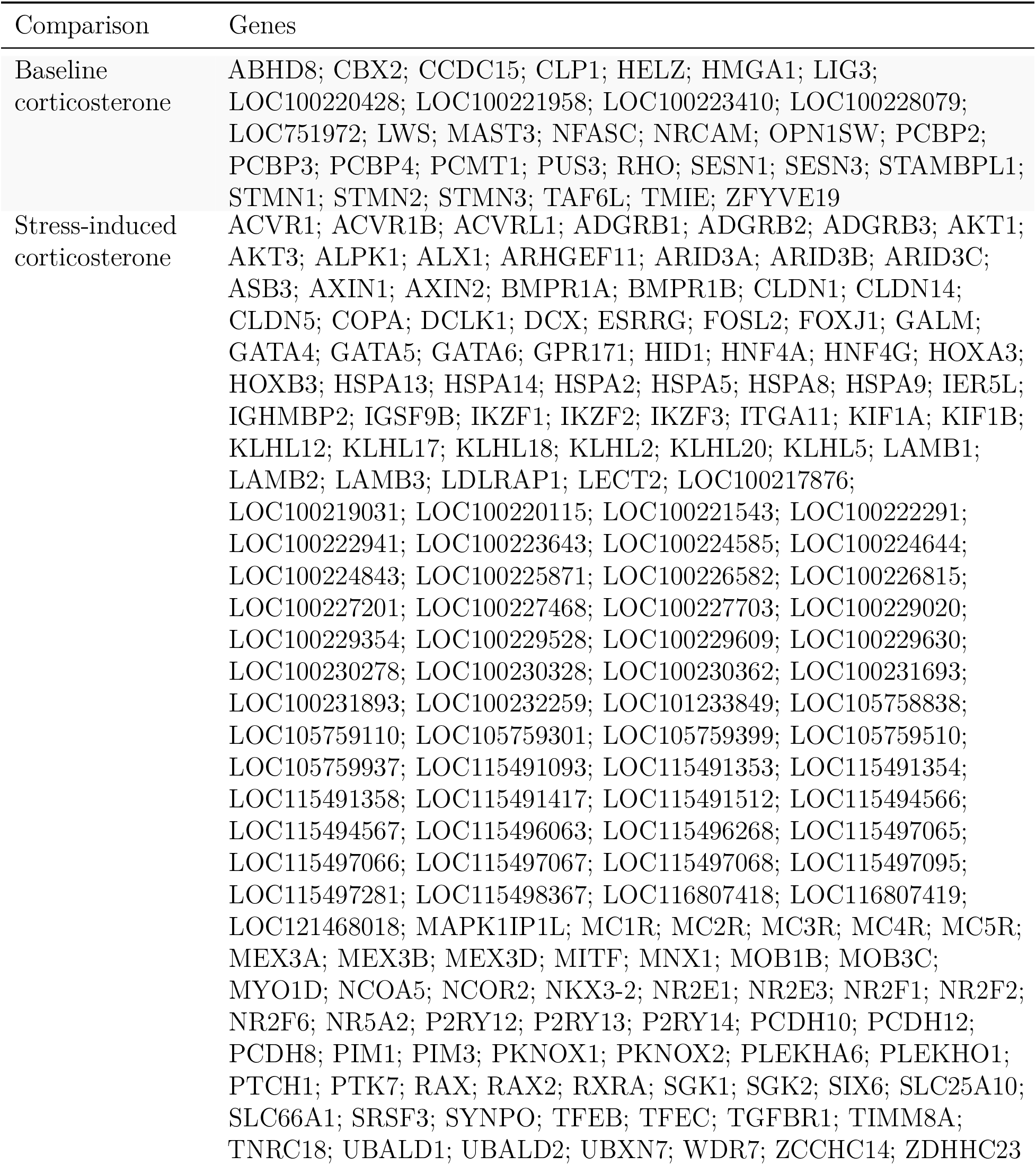
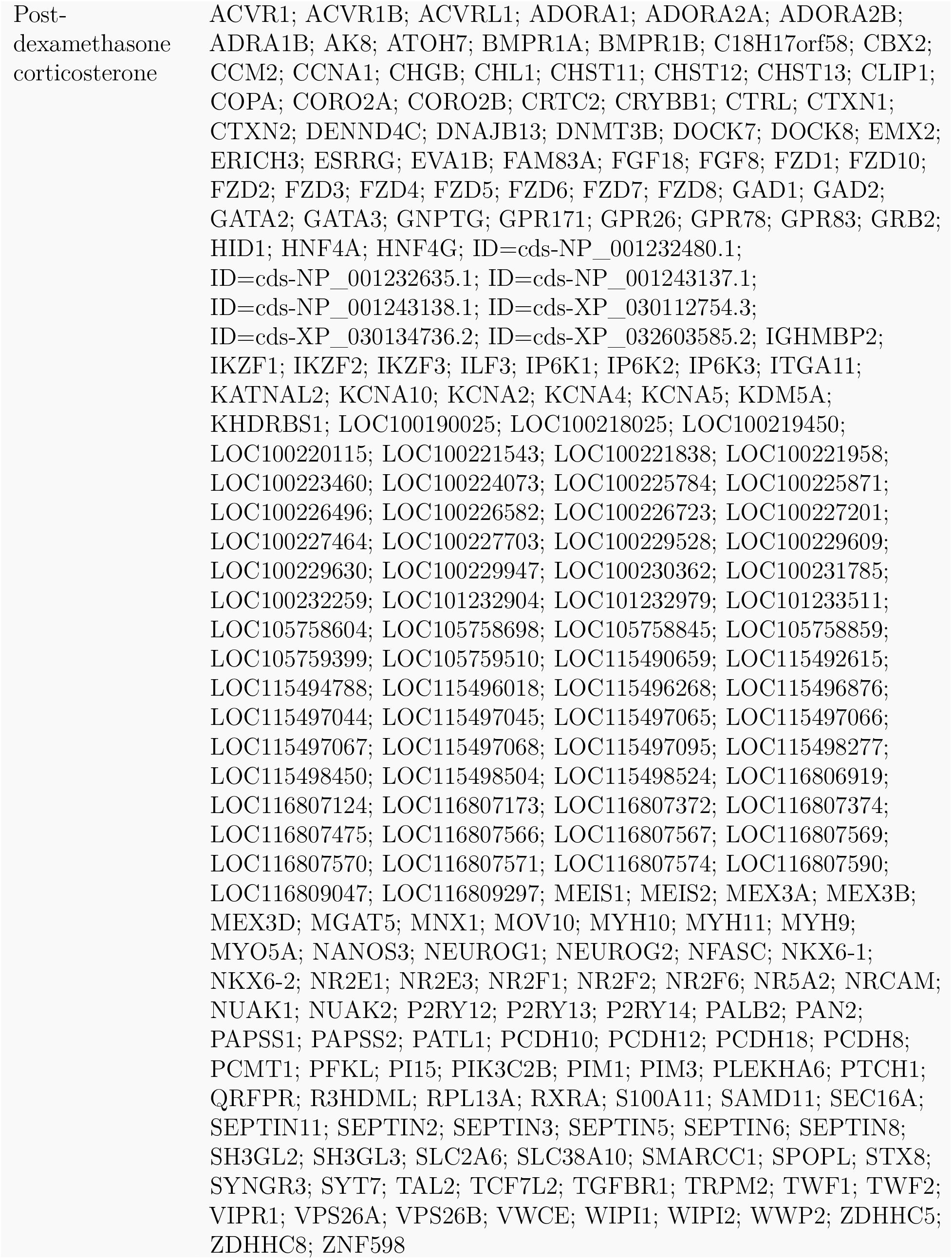
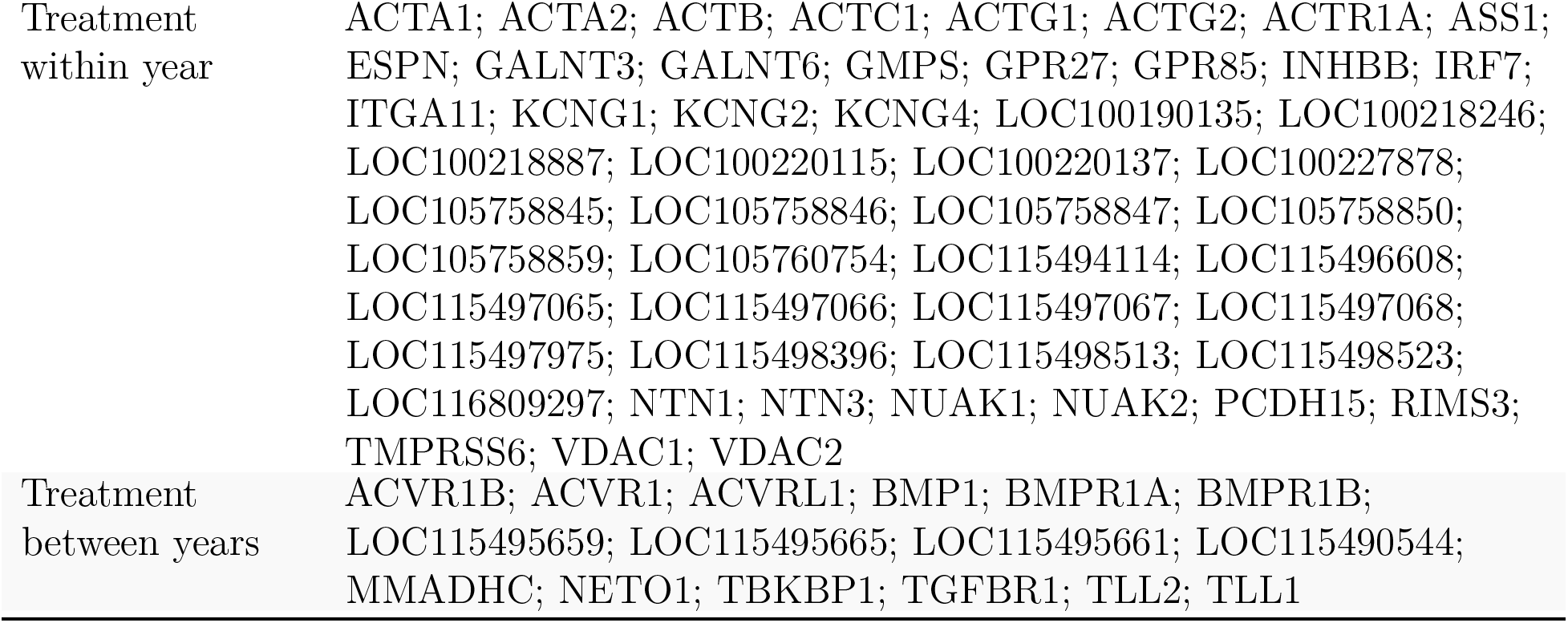
List of genes with differentially methylated CpGs in exons, introns, or within 2kb upstream of the gene.

**Table S4:** GO terms identified using DAVID after correction to a false discovery rate of 0.05 for each comparison set.

| GO Term | FDR | Function |
| --- | --- | --- |
| **Baseline corticosterone** |  |  |
| GO:0009881 | 1.23e-04 | photoreceptor activity |
| GO:0016037 | 4.43e-04 | light absorption |
| GO:0018298 | 4.43e-04 | protein-chromophore linkage |
| GO:0009583 | 4.43e-04 | detection of light stimulus |
| GO:0007602 | 4.43e-04 | phototransduction |
| GO:0009582 | 9.60e-03 | detection of abiotic stimulus |
| GO:0009581 | 9.60e-03 | detection of external stimulus |
| GO:0016038 | 9.60e-03 | absorption of visible light |
| GO:0031110 | 2.20e-03 | regulation of microtubule polymerization or depolymerization |
| GO:0051606 | 9.60e-03 | detection of stimulus |
| GO:0007601 | 3.88e-03 | visual perception |
| GO:0009605 | 1.88e-02 | response to external stimulus |
| GO:0050953 | 1.98e-02 | sensory perception of light stimulus |
| GO:0009416 | 4.79e-02 | response to light stimulus |
| **Stress-induced corticosterone** |  |  |
| GO:0004675 | 4.13e-04 | transmembrane receptor protein serine/threonine kinase activity |
| GO:0097159 | 5.86e-05 | organic cyclic compound binding |
| GO:1901363 | 6.08e-05 | heterocyclic compound binding |
| GO:0043565 | 6.12e-04 | sequence-specific DNA binding |
| GO:0003700 | 5.06e-04 | transcription factor activity, sequence-specific DNA binding |
| GO:0001071 | 9.66e-04 | nucleic acid binding transcription factor activity |
| GO:0008270 | 4.78e-03 | zinc ion binding |
| GO:0038023 | 9.15e-04 | signaling receptor activity |
| GO:0003676 | 6.34e-03 | nucleic acid binding |
| GO:0004977 | 5.48e-03 | melanocortin receptor activity |
| GO:0004674 | 5.48e-03 | protein serine/threonine kinase activity |
| GO:0004871 | 1.60e-03 | signal transducer activity |
| GO:0003677 | 3.92e-02 | DNA binding |
| GO:0005524 | 1.14e-02 | ATP binding |
| GO:0004872 | 3.85e-03 | receptor activity |
| GO:0032559 | 1.21e-02 | adenyl ribonucleotide binding |
| GO:0030554 | 1.21e-02 | adenyl nucleotide binding |
| GO:0060089 | 3.58e-03 | molecular transducer activity |
| GO:0046914 | 2.43e-02 | transition metal ion binding |
| GO:0004888 | 4.35e-02 | transmembrane signaling receptor activity |
| GO:0004930 | 4.62e-02 | G-protein coupled receptor activity |
| GO:0031625 | 3.98e-02 | ubiquitin protein ligase binding |
| **Post-dexamethasone corticosterone** |  |  |
| GO:0038023 | 2.65e-03 | signaling receptor activity |

**Table S4:** GO terms identified using DAVID after correction to a false discovery rate of 0.05 for each comparison set. (*continued*)
| GO Term | FDR | Function |
| --- | --- | --- |
| GO:0004871 | 4.04e-03 | signal transducer activity |
| GO:0042813 | 4.62e-03 | Wnt-activated receptor activity |
| GO:0004888 | 5.10e-03 | transmembrane signaling receptor activity |
| GO:0035586 | 5.10e-03 | purinergic receptor activity |
| GO:0060089 | 5.10e-03 | molecular transducer activity |
| GO:0004872 | 5.10e-03 | receptor activity |
| GO:0004675 | 8.21e-03 | transmembrane receptor protein serine/threonine kinase activity |
| GO:0099600 | 9.77e-03 | transmembrane receptor activity |
| GO:0017147 | 2.78e-02 | Wnt-protein binding |
| **Within-year treatment** |  |  |
| GO:0005200 | 1.06e-03 | structural constituent of cytoskeleton |
| GO:0005198 | 1.89e-02 | structural molecule activity |
| **Between-year treatment** |  |  |
| GO:0004675 | 1.05e-06 | transmembrane receptor protein serine/threonine kinase activity |
| GO:0004674 | 3.87e-05 | protein serine/threonine kinase activity |
| GO:0019199 | 1.62e-04 | transmembrane receptor protein kinase activity |
| GO:0004672 | 7.78e-04 | protein kinase activity |
| GO:0016772 | 1.12e-03 | transferase activity, transferring phosphorus-containing groups |
| GO:0016773 | 1.12e-03 | phosphotransferase activity, alcohol group as acceptor |
| GO:0016301 | 1.12e-03 | kinase activity |
| GO:0016740 | 3.56e-03 | transferase activity |
| GO:0004888 | 3.56e-03 | transmembrane signaling receptor activity |
| GO:0099600 | 3.56e-03 | transmembrane receptor activity |
| GO:0004871 | 2.65e-03 | signal transducer activity |
| GO:0060089 | 2.65e-03 | molecular transducer activity |
| GO:0038023 | 3.56e-03 | signaling receptor activity |
| GO:0004872 | 3.56e-03 | receptor activity |
| GO:0032559 | 4.35e-03 | adenyl ribonucleotide binding |
| GO:0030554 | 4.35e-03 | adenyl nucleotide binding |
| GO:0005524 | 4.35e-03 | ATP binding |
| GO:0003824 | 3.35e-03 | catalytic activity |
| GO:0035639 | 1.81e-02 | purine ribonucleoside triphosphate binding |
| GO:0032555 | 1.81e-02 | purine ribonucleotide binding |
| GO:0017076 | 1.81e-02 | purine nucleotide binding |
| GO:0032553 | 1.86e-02 | ribonucleotide binding |
| GO:1901265 | 1.86e-02 | nucleoside phosphate binding |
| GO:0000166 | 1.86e-02 | nucleotide binding |
| GO:0043167 | 2.60e-02 | ion binding |
| GO:0097367 | 2.80e-02 | carbohydrate derivative binding |
| GO:0036094 | 2.80e-02 | small molecule binding |

